# Elucidating the neuroanatomical correlates of social connectedness, sleep quality and psychological distress in early adolescence

**DOI:** 10.1101/2020.11.19.390336

**Authors:** Daniel Jamieson, Larisa T McLoughlin, Denise A Beaudequin, Zack Shan, Amanda Boyes, Paul Schwenn, Jim Lagopoulos, Daniel F Hermens

**Affiliations:** Thompson Institute, The Univeristy of the Sunshine Coast, Birtinya, QLD, Australia

**Keywords:** Social connectedness, Mental Health, Sleep quality, White matter, Adolescence

## Abstract

**Background:** Adolescence is an important period for developing one’s sense of self. Social connectedness has been linked to a sense of self which in turn has links to resilience in mental disorders. Adolescence is also a period of increased risk of chronic sleep deprivation during a time of ongoing white matter (WM) maturation. The complex relationship between these variables and their relationship with the onset on mental disorders during adolescence remains largely unexplored.

**Methods:** *N* = 64 participants aged 12 years (*M* = 12.6) completed the Pittsburgh Sleep Quality Index (PSQI), Social connectedness scale (SCS) and a diffusion weighted Magnetic Resonance Imaging (MRI) scan to investigate the relationship of these variables to predict psychological distress via the Kessler psychological distress scale (K10) in early adolescents. Multiple regression analysis was used with K10 entered as the dependent variable and SCS, PSQI, and values of white matter integrity as the predictor variables.

**Results:** Results showed that while all four variables collectively accounted for a significant proportion of the variance in K10 (41.1%), SCS and PSQI were the only predictors that accounted for a significant proportion of variance uniquely.

**Conclusions:** These findings suggest interventions aimed at increasing levels of social connectedness and sleep quality during adolescence may reduce psychological distress. Future longitudinal reporting of this combination of variables is suggested.

## Introduction

Social connectedness, an individual’s ability to feel comfortable, confident and a sense of belonging within a social context extending beyond family and friends [1], has been shown to be vital for good mental health and vice versa [2]. A number of studies have linked levels of social connectedness to depression and in many cases social withdrawal tends to precede a diagnosis of depression [3]. For instance, Cacioppo et al. (2010) reported that subjective perception of one’s self as socially isolated was a significant predictor of future development of a depressive disorder [4]. Being socially isolated has also been shown to reduce responsiveness to anti-depressant medication for people with major depressive disorder [5]. Moreover, a key criterion for a diagnosis of depression in the Diagnostic and Statistical Manual of Mental Disorders 5^th^ edition (DSM-5) is withdrawal from social situations as well as a lack of interest in social roles such as work or close relationships [6]. Similarly, other research has found higher levels of social connectedness decreases risk of suicidal behaviour in young people [7].

Adolescence is an important time for developing a sense of self (i.e. a person’s perception of themselves) [8] which has been shown to be strongly linked to levels of social connectedness through an increase in adolescents desire to compare themselves to their peers [8, 9]. Functional magnetic resonance imaging studies (fMRI) have demonstrated that different brain regions are activated when thinking about the self between late childhood (10 years of age) and early adulthood (26 years of age) [10], suggesting the neurobiological substrates of the sense of self are constantly developing during adolescence. Importantly, a strong sense of self has been linked to increased resilience to mental disorders [11], of which a large majority first appear during adolescence [12]. Being socially connected can reduce levels of psychological distress, depression and emotional/behavioural difficulties [13] and may result in more positive mental health and wellbeing [14].

Reduced sleep quality and mental disorders often co-occur. Mood and anxiety disorders frequently occur in tandem with sleep abnormalities [15-20] and sleep disturbance is listed as a criterion in the DSM-5 for a number of mental disorders including anxiety and depressive disorders [6]. Sleep deprivation is a major issue for many adolescents often attributed to the combination of a puberty aligned circadian rhythm phase delay [21, 22] and the need to wake early for formal activities such as early school start times [23]. This naturally occurring phase delay is often extended later into the night by light exposure via smart phone and computer tablet use which acts to further delay the release of melatonin which in turn adversely impacts sleep onset [24, 25].

A number of studies have linked levels of social connectedness with quality of sleep [26-28]. Kaushal et al. (2012) found that socially isolating mice resulted in reduced sleep quality [26] and these results have been replicated in human studies. Tavernier and Willoughby (2015) found that self-reported loneliness was associated with sleep quality with more positive social connections leading to improved emotional regulation and improved sleep [27]. Simon and Walker (2018) proposed a model where an initial lack of sleep leads to behaviour such as social withdrawal which ultimately results in loneliness. This loneliness in turn leads to poor sleep quality resulting in a self-reinforcing and detrimental cycle [28].

Adolescence is also a period of ongoing brain development which has been suggested to be influenced by experiences such as sleep [29] and social connections [30]. Brain development during adolescence involves a number of neural adaptations including myelination of white matter (WM) tracts [31]. The degree of WM myelination can be described in terms of its structural integrity [32]. Diffusion tensor imaging (DTI) is commonly utilised to examine the structural integrity of WM tracts [33]. From DTI, fractional anisotropy (FA) a directional measure of the diffusion of water within and along WM tracts can be determined [34]. FA values are commonly reported in studies that focus on the structural integrity of WM tracts [33].

Previous studies have demonstrated associations between grey matter volumes (GMV) and the size of an individual’s social group [35-38], however, WM development has largely been neglected. One study involving WM conducted by Szeligo and Leblond (1977) found that rats raised in an enriched environment showed increased levels of myelination compared to those raised in impoverished environments [39]. In humans, Hampton et al. (2016) found evidence to suggest that WM connectivity between the amygdala, orbitofrontal cortex and anterior temporal lobe may be associated with the size of a person’s social network [40].

In a recent study by our group, WM microstructure of the posterior limb of the internal capsule (PLIC) was shown to be significantly correlated with self-reported sleep quality [41]. In a study of early adolescents, those who reported poorer sleep quality were shown to have higher FA values of the posterior limb of the internal capsule (PLIC) along with a similar although non-significant trend for the anterior limb of the internal capsule (ALIC) [41]. Higher FA value of the PLIC has also been found to be positively correlated with fear of anxiety-related symptoms in a study of patients with panic disorder [42]. While increased PLIC FA value has been linked to major depressive disorder (MDD) with a recent study showing greater FA value of the PLIC in a MDD group compared to controls [43]. Combined, these findings suggest further investigation of the role the PLIC and ALIC in adolescent psychological distress is necessary along with an investigation into any possible relationship with social connectedness.

In the current study, participants aged 12 years (*M* =12.6, *SD* .32) completed the Social Connectedness Scale (SCS), Pittsburgh Sleep Quality Index (PSQI), Kessler Psychological distress scale (K10) and diffusion weighted MRI to investigate the collective and individual influence of social connectedness, self-report sleep quality, and WM microstructure of the PLIC and ALIC on psychological distress. It was hypothesised that: i) a regression model consisting of SCS, PSQI and the FA value of the PLIC and ALIC will predict a significant proportion of the variance in K10 scores in a sample of early adolescents, and ii) that scores on the SCS, PSQI, and FA values of the ALIC and PLIC would all make significant contributions to this regression model.

## Methods

### Ethical Approval

Ethics approval was granted by the USC Human Research Ethics Committee as part of the Longitudinal Adolescent Brain Study (LABS) (Approval Number: A181064). Informed assent and consent was obtained from all participants and their guardian/s.

### Participants

Participant data were accessed from the Longitudinal Adolescent Brain Study (LABS) being undertaken at the Thompson Institute, USC. Inclusion criteria for LABS was that at study entry all participants were 12 years of age and in grade 7 (first year of secondary school). Participants were recruited from the Sunshine Coast region and were proficient in spoken and written English. Participants who reported suffering from a major neurological disorder, intellectual disability, major medical illness or who reported having sustained a head injury that involved loss of consciousness for greater than 30 minutes were excluded.

### Measures

#### Sleep Quality

Participants undertook the Pittsburgh Sleep Quality Index (PSQI), an 18-item self-report, retrospective (past month) questionnaire [44]. The PSQI is designed to measure seven components of sleep: subjective sleep quality, sleep latency, sleep duration, habitual sleep efficiency, sleep disturbance, use of sleep medication, and daytime dysfunction. Responses are provided on a four-point Likert scale and tallied to give an overall score. Subscale scores are rank-ordinal in nature, however, the PSQI total score, which is calculated by the summation of all seven subscale scores, can be treated as a continuous variable with potential scores ranging from 0 to 21, with lower scores indicating better sleep quality. Buysse et al. (1989) suggest that scores greater than 5 are associated with poor sleep quality [44]. Although the PSQI was initially validated using an adult sample [44] it has since been shown to be a reliable and valid measure of sleep quality and quantity for use in a wide range of populations including adolescents and young adults [45, 46].

#### Psychological Distress

Participants also completed the Kessler psychological distress scale (K10) [47], a 10-item self-report questionnaire designed to measure depression and anxiety levels on a five-point Likert scale by asking about feelings over the past 30 days. Individual item scores are then tallied to provide an overall score ranging from 10 to 50 with higher scores indicating increased psychological distress. The K10 has been shown to have good reliability and validity across cultures and clinical populations, and is used as a proxy of mental disorder at various levels of severity [48-50].

#### Social Connectedness

As part of the self-report questionnaire, participants completed a revised 15 item version of the Social Connectedness Scale (SCS) [51]. Participants respond on a six-point Likert scale ranging from 1 = *Strongly Disagree* to 6 = *Strongly Agree*. Internal reliability for the SCS is excellent, with a Cronbach’s alpha coefficient of .92 [52]. The SCS demonstrates appropriate convergent and discriminant validity [52]. The SCS captures elements of closeness to others, sense of togetherness and connection [52].

#### MRI acquisition

All participants also underwent a MRI protocol at the Thompson Institute. All scans were acquired on a 3 Tesla Siemens Skyra (Erlangen, Germany) scanner using a 64-channel head and neck coil. Structural MRI was acquired using a Magnetization Prepared Rapid Acquisition Gradient Echo (MP-RAGE) T1-weighted sequence (TR=2200ms, TE=1.76/850ms, Resolution= 0.9/0.9/0.9, FOV 240mm). Diffusion weighted imaging (DWI) was acquired using a single shot spin-echo EPI technique, optimised for minimum error within the tensor measurement, enabling the measurement of white matter microstructure. It included a 96-direction DWI optimised for determining crossing fibres (56 slices, no slice gap, slice thickness = 2.0mm, in-plane resolution = 2×2mm; TR=3300 msec, TE=115msec, FOV 228 mm, 3 B-values or “shells” were acquired at 30 at b = 1000 s/mm2, 60 at b=2500s/mm2 and 6 at b=0 repetitions, duration= 9 minutes). A reversed phase encoding “blip down sequence with 6 b=0 repetitions was also be acquired for EPI distortion correction during post processing.

#### Image Processing

Preprocessing of raw images was carried out using FSL [53]. B0 images were extracted in the AP and PA directions and merged using fslroi and fslmerge tools followed by susceptibility distortion correction using the fsl topup tool [54] and distortion correction using eddy [55]. Voxelwise statistical analysis of the FA data was carried out using TBSS (Tract-Based Spatial Statistics) [56], part of FSL [53]. First, FA images were created by fitting a tensor model to the raw diffusion data using FDT, and then brain-extracted using BET [57]. All FA data were then aligned into common space using the nonlinear registration tool FNIRT, which uses a b-spline representation of the registration warp field [58]. Then, the mean FA image was created and thinned to create a mean FA skeleton which represents the centres of all tracts common to the group. Each subject’s aligned FA data was then projected onto this skeleton and the resulting data fed into voxelwise cross-subject statistics.

The JHU WM tractography atlas and JHU ICBM DTI 81 labels atlas [59] (Part of FSL) were used to create tract specific masks of the tracts of interest and FSLMATHS and FSL MEANTS commands were used to extract the tract specific FA values for statistical analysis [53].

#### Statistical Analysis

SPSS® Version 26 (SPSS Inc., Chicago, Illinois, USA) was used to produce sample descriptive statistics. A standard multiple regression analysis was then performed with K10 entered as the dependent variable and SCS, PSQI score and FA values of the ALIC and PLIC entered as the predictor variables. Regression assumptions were deemed as met following visual inspection of the normal P-P plot and scatterplot of the residuals and VIF statistics were all under 3 (less than 5 is considered good).

## Results

Table 1 provides the sample descriptive statistics *N*=64 (33 Female).

**Table 1.** Descriptive statistics for the sample of early adolescents.

|  | Min | Max | Mean | *SD* |
| --- | --- | --- | --- | --- |
| Age (years) | 12.1 | 13.1 | 12.6 | 0.32 |
| PSQI | 0 | 14 | 4.73 | 2.69 |
| K10 | 10 | 31 | 15.36 | 4.58 |
| SCS | 24 | 89 | 73.19 | 14.12 |
| PLIC (FA) | 0.689577 | 0.785859 | 0.730843 | 0.023 |
| ALIC (FA) | 0.537726 | 0.650816 | 0.607561 | 0.025 |

The Model summary R square statistic showed that the regression model explained 41.1% of the variance in K10 scores and the ANOVA output of the multiple regression analysis showed that this proportion of the variance was significant *F*(4,45) = 7.85, *p<*.001. Table 2 shows that the contribution of each independent variable to the overall regression model. The multiple regression revealed that social connectedness (*p =* .011) and PSQI score (*p =* .006) contributed significantly to the regression model while the FA value of the PLIC (*p =* .591) and ALIC (*p =* .334) did not make significant contributions.

**Table 2.** Findings for each independent variable

|  | B | STD Error | *t* | Sig. | Tolerance | VIF |
| --- | --- | --- | --- | --- | --- | --- |
| Constant | 23.64 | 17.51 | 1.351 | .184 |  |  |
| SCS | -.102 | .039 | -2.638 | .011* | .666 | 1.502 |
| PSQI | .569 | .199 | 2.857 | .006** | .763 | 1.311 |
| PLIC | 18.399 | 33.993 | .541 | .591 | .334 | 2.995 |
| ALIC | -28.049 | 28.745 | -.976 | .334 | .381 | 2.626 |

\*Significant at $p < .05$
\*\*Significant at $p < .01$

## Discussion

Adolescence is an important time for the development of social connections. It is a sensitive period for the development of the neural substrates of the social brain [60] and for brain development in general [32, 61, 62]. It is also a period where the majority of mental health disorders first emerge [12] and a period of heightened risk of chronic sleep deprivation [15, 18, 21-23]. It’s therefore not surprising that independently, social connectedness and sleep quality have been shown to be important for mental health [2, 15, 16, 18-20]. Few studies however have investigated the relationship between these factors in a sample of young adolescents.

The current study was designed to investigate the influence of social connectedness, sleep quality, and WM structural integrity of the ALIC and PLIC on psychological distress. It was hypothesised that firstly, a regression model consisting of SCS, PSQI and the structural integrity of the PLIC and ALIC would predict a significant proportion of the variance in K10 scores in a sample of early adolescents. Secondly it was hypothesised that scores on the SCS, PSQI, and structural integrity of the ALIC and PLIC will all make significant contributions this regression model. These hypotheses were based on previous findings showing associations between social connectedness and mental health [2-5, 7], the importance of adolescence as a period for developing a sense of self [8, 9], ongoing brain development during adolescence [10], sleep quality and mental health [15-19], the increased likelihood of poor sleep during adolescence [21, 22], social connectedness and sleep quality [26-28] and sleep quality and WM microstructure of the PLIC and ALIC [41].

In accordance with our first hypothesis, the overall model predicted a significant proportion of the variance in K10 scores. Collectively the predictors included in the model explained a significant 41.1% of the variance in K10 score in our sample of early adolescents.

With regards to the second hypothesis, SCS and PSQI both explained significant proportions of the variance in K10 scores. However, FA values of the PLIC and ALIC did not.

The findings from the current study suggest that social connectedness and sleep quality are both important predictors of psychological distress and should be areas of focus for psychological interventions during early adolescence. Currently the best practice for child and adolescent anxiety and depressive disorders based on evidence-based findings is cognitive behaviour therapy (CBT) [63], some studies have also demonstrated the effectiveness of CBT for improving sleep quality [64]. A randomised control study conducted by de Bruin et al. (2015) reported that participants in both an internet therapy group and group therapy participants showed improved sleep efficiency, sleep onset latency, and total sleep time improvements compared with waiting list controls [64]. The findings reported by de Bruin et al. (2015) demonstrate the ability to improve sleep using CBT based therapies, which results from the current study suggest may be likely to have positive psychological benefits.

Social connectedness has been shown to impact resilience in a variety of mental disorders and their related symptoms, including lower levels of depression, reduced suicide ideation, reduced levels of conduct disorder, as well as increased levels of self-esteem [65]. Interventions that increase social connectedness have been shown to be successful in improving the mental health of participants including depression, anxiety, bipolar disorder and schizophrenia [66].

The present findings suggest that the impact of social connectedness and sleep quality on psychological distress occur independently of WM maturation of the PLIC or ALIC. These two WM tracts were included in the current study based on previous findings linking these tracts to reduced sleep quality in adolescence [41]. Future studies need to broaden the search for neuroanatomical links between social connectedness, sleep and psychological distress by investigating volumetric measures of WM, as well as looking at relationships between the volume of sub-cortical grey matter structures and cortical thickness measures. One region in particular is the medial prefrontal cortex (mPFC) which has been shown to play an important role in the adolescent ‘social brain’ [60]. The mPFC has been shown to be more active in adolescents than adults during mentalising tasks which are suggested to be important for social interactions [60]. The connectivity between the mPFC and amygdala has also been shown to be associated with the regulation of emotions linking the mPFC to possible symptoms of mental disorders [67].

Some limitations of the current study need to be acknowledged. Firstly, the use of a self-report measure for sleep quality is a limitation of the current study. The PSQI was chosen due to its demonstrated validity and reliability in studies of sleep quality of adolescents [45, 46]. Importantly the PSQI has demonstrated convergent validity with measures of depression and anxiety in an adolescent sample [46]. It is acknowledged that future replication of this study would benefit from the inclusion of an objective measure of sleep quality such as polysomnography or actigraphy to provide an objective measure of sleep quality.

Secondly, reporting data collected from a single timepoint has its limitations as it only provides a snapshot of what these relationships look like. Future reporting of longitudinal data from multiple timepoints will allow investigation of patterns of relationships between variables which will assist in making better informed inferences of relationships.

## Conclusion

In conclusion, our study builds on an emerging literature by reporting findings on the associations between social connectedness, sleep quality, PLIC and ALIC WM microstructure and psychological distress. The results of our study indicate that a diagnostic model based on measures of social connectedness and self-report sleep quality may be useful for predicting levels of psychological distress in early adolescents. These findings suggest interventions aimed at increasing levels of social connectedness and sleep quality during adolescence may be important inclusions in a treatment program for adolescents with heightened psychological distress.

## Acknowledgments

This research is supported by an Australian Government Research Training Program (RTP) Scholarship and a grant from the Australian Commonwealth Government’s ‘Prioritizing Mental Health Initiative’ (2018-19).

